# Probability of Lateral Instability While Walking on Winding Paths

**DOI:** 10.1101/2024.07.16.603791

**Authors:** Anna C. Render, Joseph P. Cusumano, Jonathan B. Dingwell

## Abstract

People with balance impairments often struggle performing turns or lateral maneuvers, which can increase risk of falls and injuries. Here we asked how people’s mediolateral balance is impacted when walking on non-straight winding paths. Twenty-four healthy adults (12F/12M; 25.8±3.5 yrs) participated. Each walked on each of six paths projected onto a treadmill, comprised of three pseudo-random path oscillation frequency combinations (straight, slowly-winding, quickly-winding), each presented at either wide or narrow width. We quantified stepping errors as the percent of steps taken off each path. We quantified minimum mediolateral Margin of Stability (*MoS*_*L*_) at each step and calculated means (*μ*) and standard deviations (*σ*) for each trial. We calculated lateral Probability of Instability (*PoI*_*L*_) as participants’ statistical risk of taking unstable (*MoS*_*L*_ < 0) steps. On *narrower* paths, participants made more stepping errors and walked with smaller *μ*(*MoS*_*L*_) on all paths (p < 0.001), and exhibited increased *PoI*_*L*_ on the straight and slowly-winding paths (p < 0.001). On *winding* paths, participants made progressively more stepping errors and walked with smaller *μ*(*MoS*_*L*_) for increasingly sinuous narrow paths (all p < 0.001) and wide quickly-winding paths (all p < 0.001). They also consistently walked with larger *σ*(*MoS*_*L*_), and increased *PoI*_*L*_ on higher sinuosity paths of both widths (all p < 0.001). Though many took numerous unstable steps, no participant *fell*. Our results demonstrate healthy adults’ ability both to trade off increased risk of lateral instability for greater maneuverability, and to employ highly-versatile stepping strategies to maintain balance while walking.

## INTRODUCTION

Maneuvering constitutes a significant proportion of daily activities (Glaister et al., 2007). Walking around corners (Bergsma et al., 2021; Tillman et al., 2022), in crowded environments (Degond et al., 2013), through narrow spaces, or negotiating obstacles (Musselman and Yang, 2007) are fundamental to our mobility. However, executing maneuvers imposes that we adapt our stepping (Desmet et al., 2022; Hak et al., 2013; Ochs et al., 2021; Twardzik et al., 2019). Such tasks may heighten our risk of instability (Bruijn and van Dieën, 2018; Desmet et al., 2022; Desmet et al., 2024), particularly as people age and face challenges in weight shifting (Patla et al., 1993) and balance control (van Dieën et al., 2005). Indeed, older adults often fall when trying to perform such lateral maneuvers (Robinovitch et al., 2013). This can pose potential for serious injury (Parkkari et al., 1999; Yang et al., 2020). Thus, it is important to better understand how executing lateral maneuvers affects mediolateral balance while walking.

One metric widely used to quantify walking balance is the lateral margin of stability (*MoS*_*L*_) (Hof, 2008). *MoS*_*L*_ quantifies a *margin* as the linear distance from a *threshold* (i.e., *MoS*_*L*_ = 0) beyond which stability is presumed to be compromised (Hof et al., 2005; Kazanski et al., 2024). However, in multiple *de*stabilizing contexts, people exhibit larger (not smaller) mean values of *MoS*_*L*_ (Hak et al., 2012; Hurt and Grabiner, 2015; Onushko et al., 2019; Wu et al., 2017). Such larger mean *MoS*_*L*_ are often interpreted to indicate greater stability (Watson et al., 2021), which is quite counterintuitive. This arises because interpreting mean *MoS*_*L*_ values as indicating stability runs contrary to *MoS*_*L*_’s definition as a *margin*. We proposed a different statistic, lateral Probability of Instability (*PoI*_*L*_) (Kazanski et al., 2022), that is computed from the same *MoS*_*L*_ values as the mean, but characterizes an attribute of the distribution consistent with Hof’s original definition. *PoI*_*L*_ resolves the paradox of misinterpreting mean *MoS*_*L*_ (Kazanski et al., 2022), and explains why perturbed/impaired people take wider steps to maintain balance (Dean et al., 2007). When people can readily take wider steps, doing so will *increase* their mean *MoS*_*L*_. But when this offsets greater step-to-step variability of *MoS*_*L*_ (McAndrew Young et al., 2012; Onushko et al., 2019), it can simultaneously *decrease PoI*_*L*_ (Kazanski et al., 2024).

However, most often when people walk, they follow paths with lateral boundaries that restrict where they can step (Dingwell and Cusumano, 2019; Kazanski et al., 2023). These paths sometimes have well-defined boundaries, like hallways, sidewalks, etc., or sometimes less-well-defined boundaries, like outdoor walking trails (Matthis et al., 2018), etc. Such lateral path boundaries limit people’s stepping options for maintaining balance. Thus, when people walk on narrow paths or beams, they exhibit smaller mean *MoS*_*L*_ (Arvin et al., 2016; da Silva Costa et al., 2020; Kazanski et al., 2024; Schrager et al., 2008).

Similarly, when people make discrete turns or maneuvers, *MoS*_*L*_ typically decreases on so-called “inside” steps (where the stepping foot is ipsilateral to direction of the maneuver) and increases on so-called “outside” steps (where the stepping foot is contralateral to the direction of the maneuver) (Acasio et al., 2017; He et al., 2018; Ho et al., 2023; Wu et al., 2015). The smaller *MoS*_*L*_ on inside steps implies that people are more unbalanced in the direction of the maneuver. This, however, simultaneously facilitates their ability to execute that maneuver (Desmet et al., 2022). Hence, people trade-off compromising their stability to enhance their lateral maneuverability (Acasio et al., 2017; Desmet et al., 2022).

Real-world paths are often both laterally constrained *and* non-straight (Bergsma et al., 2021; Matthis et al., 2018; Moussaïd et al., 2011). Experiments have not yet assessed how these competing factors combine to affect *MoS*_*L*_. Therefore, to simulate real-world-like walking conditions that impose continuously-changing balance challenges, here we created winding walking paths. We investigated the influence of these paths’ lateral oscillation frequency, or sinuosity, by creating paths with either slowly-winding or quickly-winding curves. We also tested the influence of path width by creating wide and narrow versions of each winding path.

We hypothesized that people walking on narrow paths would take more unstable steps due to the constraints placed on foot placement. We hypothesized that people walking on windier (i.e., higher sinuosity) paths would likewise take more unstable steps due to their continuously changing direction. Lastly, we hypothesized that each of these two factors (path width and sinuosity) would compound each other, such that people would take even more unstable steps when both challenges were imposed simultaneously.

By investigating how varying degrees of continuous maneuvering affect one’s probability of experiencing lateral balance loss, we further and better characterize how people make trade-offs between stability and maneuverability (Acasio et al., 2017; Desmet et al., 2022; Wu et al., 2017). This research holds particular significance for high fall-risk populations like older adults (Robinovitch et al., 2013), as it offers insights into identifying everyday tasks that may impose a challenge to their stability. By quantifying risk of lateral instability, this work both expands our understanding of human locomotion and contributes to developing strategies aimed at enhancing mobility and mitigating fall risk in diverse populations.

## METHODS

### Data Availability

All relevant data underlying the results reported here are available on Dryad ([Dataset] Render et al., 2024).

### Participants

Prior to participation, twenty-four participants (Table 1) provided written informed consent, as approved by the Institutional Review Board of Penn State University. Participants were screened to ensure no medications, lower limb injuries, surgeries, musculoskeletal, cardiovascular, neurological, or visual conditions affected their gait.

**Table 1.** Participant characteristics. All values except Sex are given as Mean ± Standard Deviation.

| Characteristic: | Value: |
| --- | --- |
| Sex [F/M] | 12 / 12 |
| Age [yrs] | 25.77 ± 3.53 |
| Body Height [m] | 1.74 ± 0.11 |
| Body Mass [kg] | 70.15 ± 14.75 |
| Body Mass Index [kg/m<sup>2</sup>] | 22.97 ± 3.10 |
| Leg Length [m] | 0.93 ± 0.05 |

### Experimental Protocol

Participants walked on a 1.2 m wide treadmill in an M-Gait virtual reality system (Motek Medical, Netherlands; Fig.1A). Each participant wore a safety harness. All walking trials were performed at 1.2 m/s. Participants first walked 4-minutes to acclimate to the system.

Participants were asked to walk on paths projected onto the treadmill belt (Fig. 1A). Five meters (5 m) of moving path were projected at all times to allow participants more-than-sufficient visual information to plan their steps (Matthis et al., 2017; Matthis et al., 2018). We created these paths using a sum of sin waves:

**Figure 1.**
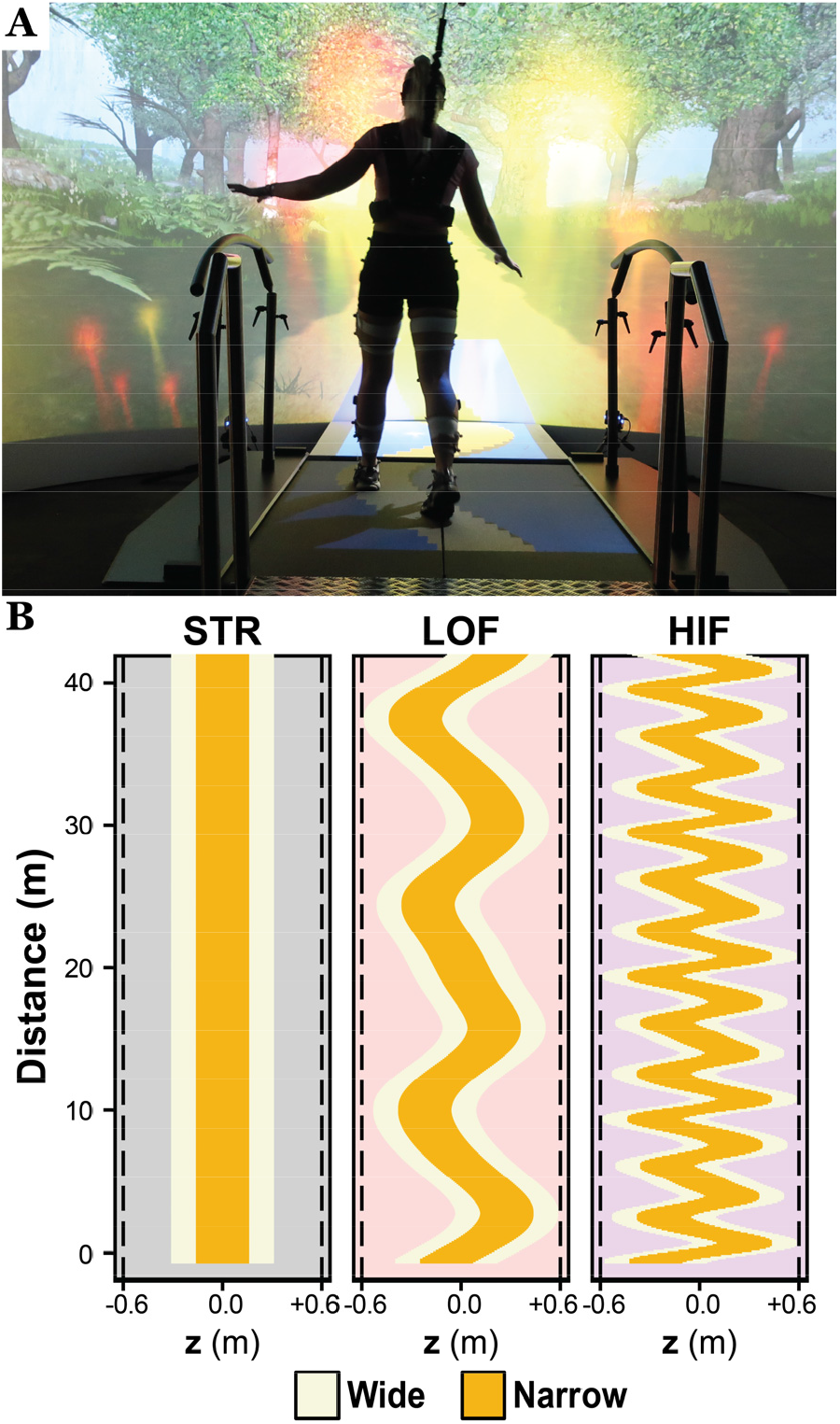
**A)** Photo of a participant walking in the Motek M-Gait virtual reality system with a path projected onto the treadmill surface. **B)** Participants traversed paths of each of three oscillation frequency combinations (Eq. 1): straight (STR; grey panel), low frequency (LOF; red panel), and high frequency (HIF; purple panel). Each path frequency was presented at each of two different path widths, wide (W = 0.6m; beige) and narrow (N = 0.3m; yellow), for a total of six unique conditions.

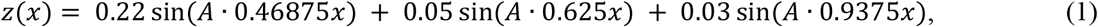

where *z* is the lateral position (in meters) of the path center, *x* is forward distance (in meters), and A is a frequency scaling factor. Each participant walked on each of six unique paths (Fig. 1B), combining either of two path widths, wide (W = 0.60 m) and narrow (N = 0.30 m), with each of three oscillation frequencies: straight (STR: *A* = 0), low frequency (LOF: *A* = 1), and high frequency (HIF: *A* = 4).

We instructed participants to “stay on the path” and minimize stepping errors. They received visual (firework) and auditory (flute) penalties when steps landed outside the path boundaries. For each condition, participants completed a 1-minute introductory trial followed by two 4-minute experimental trials. Order of presentation of conditions was randomly assigned to each participant and counterbalanced across participants using a Latin Square design. Participants were allowed to rest as needed after each trial.

### Data Processing

Each participant wore 16 retroreflective markers: four around the head, four around the pelvis (left and right PSIS and ASIS), and four on each shoe (first and fifth metatarsal heads, lateral malleolus, and calcaneus). Kinematic data were collected at 100 Hz from a 10-camera Vicon system (Oxford Metrics, Oxford, UK) and post-processed using Vicon Nexus software. Marker trajectories and path data from D-Flow software (Motek Medical, Netherlands) were analyzed in MATLAB (MathWorks, Natick, MA).

For each experimental trial, we low-pass filtered (10 Hz 4^th^-order Butterworth) the raw marker trajectories and interpolated these data to 600 Hz (Bohnsack-McLagan et al., 2016). Heel strikes and toe-offs were identified using a previously validated algorithm (Zeni et al., 2008). For consistency, we analyzed the first *N* = 350 steps of each trial.

On a curved path, one’s direction of motion changes at each new step. We therefore transformed our data into path-based local coordinates (Dingwell et al., 2023; Ho et al., 2023). At each step, we aligned this path-based local coordinate system to be tangent to the point on the path closest to the midpoint between the left and right foot placements for that step (Dingwell et al., 2023). The locally lateral (*z*-axis) motion was then taken to be perpendicular to the path at that point on the path.

### Stepping Errors

The instructed task objective was to remain on the path. We counted as a stepping error any step where any of the first or fifth metatarsal head or calcaneus markers exceeded the path boundary. We then quantified Stepping Errors as the percentage of steps (out of *N* = 350) within each trial that landed outside the path’s boundaries. This measure served as a functional measure of overall task performance.

### Lateral Margins of Stability

We quantified lateral stability in the locally-lateral *z*-direction at each step (Fig. 2A). We took the average motion of the 4 pelvis markers to approximate the center-of-mass (CoM) state (*z, ż*) (Havens et al., 2018). The extrapolated center-of-mass (*XCoM*) position was then (Hof et al., 2005):

**Figure 2.**
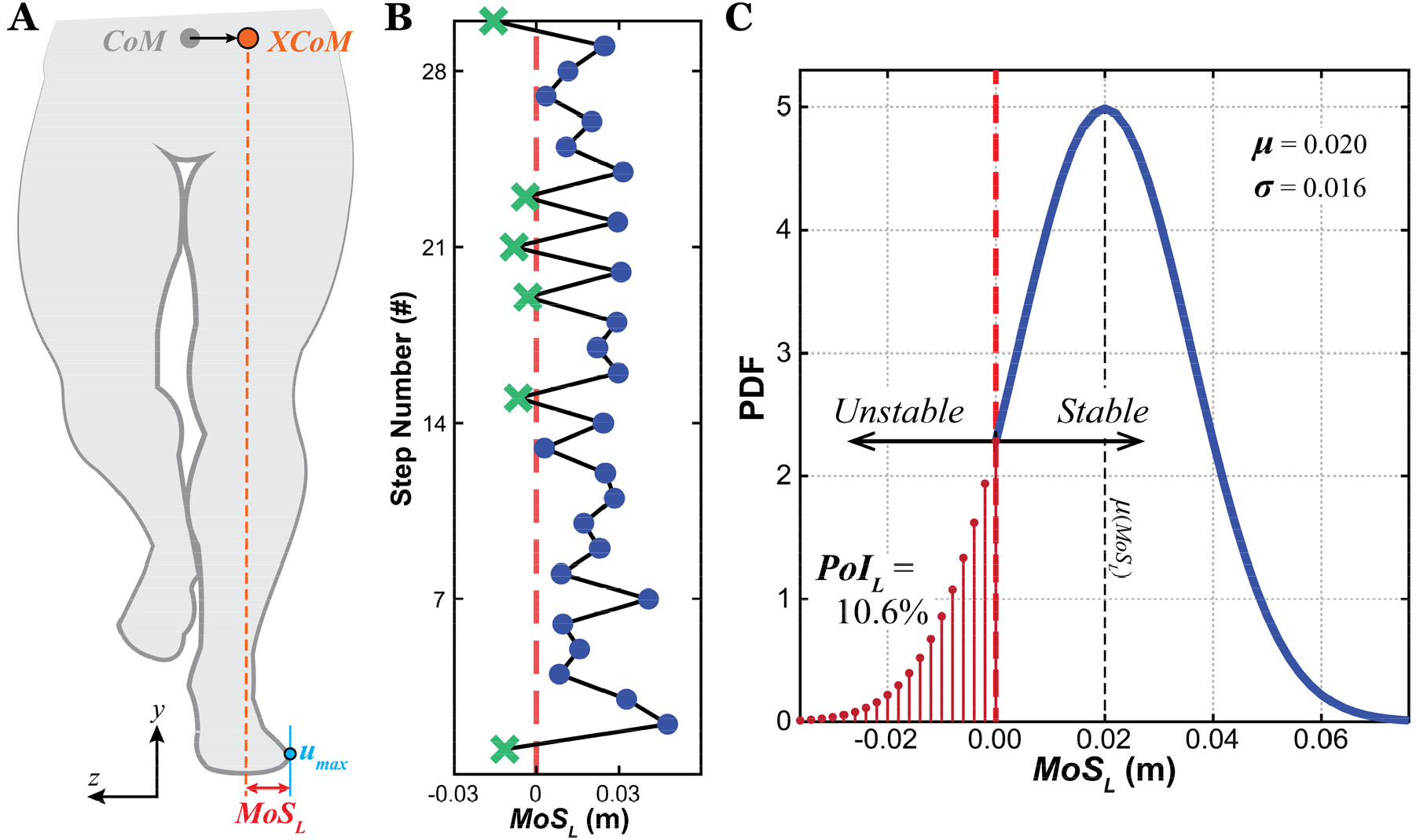
**A**) Schematic showing the relevant variables extracted at each step to calculate *MoS*_*Lmin*_ (Eq. 2-3): center-of-mass (CoM; grey), extrapolated center-of-mass (*XCoM*_*L*_; orange), and lateral base-of-support boundary (*u*_*max*_; blue). **B**) Example time series data of *MoS*_*Lmin*_ across several steps. The vertical red dashed line indicates *MoS*_*Lmin*_ = 0. Values of *MoS*_*Lmin*_ > 0 (blue • markers) are considered stable, while values of *MoS*_*Lmin*_ < 0 (green × markers) are considered unstable. **C**) Hypothetical probability density function (PDF) to demonstrate how to calculate *PoI*_*L*_ from a distribution of *MoS*_*Lmin*_ values (e.g., as might be obtained from **B**, etc.). Here, *μ*(*MoS*_*Lmin*_) = 0.020 and *σ*(*MoS*_*Lmin*_) = 0.016. The dashed vertical red line indicates the stability threshold separating stable (*MoS*_*Lmin*_ > 0) from unstable steps (*MoS*_*Lmin*_ < 0). Note that the mean, *μ*(*MoS*_*Lmin*_) = 0.020, is well within the *stable* region, and so yields no information regarding *un*stable steps. Conversely, *PoI*_*L*_ (Eq. 4) directly calculates the cumulative probability (area under the curve) precisely of those *un*stable (*MoS*_*Lmin*_ < 0) steps. Here *PoI*_*L*_ = 10.6%, which equates to taking 1 unstable step every ∼9.4 steps. Thus, *PoI*_*L*_ directly estimates statistical *risk* of instability, consistent with Hof’s theory, while *μ*(*MoS*_*Ln*_) does not.

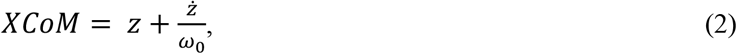

where 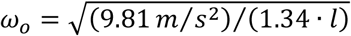 and *l* was leg length, as measured from the greater trochanter to the lateral malleolus (Curtze et al., 2024). We took the *z*-position of the leading foot’s fifth metatarsal-phalangeal joint marker to approximate the lateral boundary of the base of support (*u*_*max*_). We calculated the *minimum MoS*_*L*_ within each step *n* as:

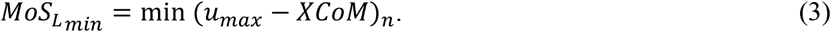

Hence, a positive *MoS*_*Lmin*_ indicated a stable step, while a negative *MoS*_*Lmin*_ indicated an unstable step.

For each trial performed by each participant for each condition, we extracted time series of *MoS*_*Lmin*_ (e.g., Fig. 2B) from all *n* ∈ {1, …, 350} steps. We then computed the within-trial means, *μ*(*MoS*_*Lmin*_), and standard deviations, *σ*(*MoS*_*Lmin*_), for each trial.

### Lateral Probability of Instability

As defined (Hof et al., 2005), *MoS*_*L*_ is a *margin* that quantifies the distance to a defined *threshold* for becoming unstable (i.e., *MoS*_*L*_ = 0). While researchers commonly average *MoS*_*L*_ across steps in a trial (Watson et al., 2021), this is *not* consistent with how *MoS*_*L*_ was originally defined (Kazanski et al., 2022, 2024), because *μ*(*MoS*_*L*_) does not quantify a person’s risk of exceeding that threshold (i.e., of experiencing *MoS*_*L*_ < 0) and thus becoming unstable.

Therefore, for each trial, we estimated the lateral *Probability of Instability* (*PoI*_*L*_) (Kazanski et al., 2022) as the cumulative probability (P), for an assumed normal distribution of *MoS*_*Lmin*_, that steps within that trial would exceed the *MoS*_*Lmin*_ < 0 threshold (Fig. 2C):

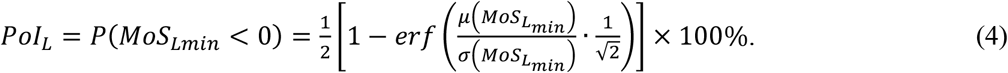

For example, a *PoI*_*L*_ = 10% implies that a person is likely to take ∼35 unstable (*MoS*_*Lmin*_ < 0) steps over a 350 step trial. Importantly, *PoI*_*L*_ is neither “new” or “different” from *MoS*_*L*_: it simply extracts a different statistical metric than the mean from the exact same distribution of *MoS*_*Lmin*_ values (Fig. 2B-C).

### Statistical Analyses

We applied a two-factor (Frequency × Width) mixed-effects analyses of variance (ANOVA) with repeated measures to test for differences between conditions for each dependent measure: % Stepping Errors, *μ*(*MoS*_*Lmin*_), *σ*(*MoS*_*Lmin*_) and *PoI*_*L*_. For *σ*(*MoS*_*Lmin*_), to satisfy normality assumptions, we first log-transformed these data. For % Stepping Errors and *PoI*_*L*_, because multiple trials yielded resulting values equal to 0, we first added a small constant value (0.001) before we log-transformed these measures. When we found main or interaction effects to be significant, we performed Tukey’s pairwise post-hoc comparisons to test for specific differences between path frequencies (STR, LOF, HIF) for each path width (W, N), and between path widths for each path frequency. All statistical analyses were conducted using Minitab (Minitab, Inc., State College, PA).

## RESULTS

### Stepping Errors

Participants took more steps off the narrower paths than the wide paths for all path oscillation frequencies (p < 0.001; Fig. 3; Table 2). Similarly, participants made more stepping errors as path oscillation frequency increased (from STR to LOF to HIF) for all narrow paths (all p ≤ 0.007). However, for the wide paths, they only made more stepping errors on the HIF paths (each p < 0.001) (Fig. 3; Table 2).

**Table 2.** Statistical results for differences between path Frequencies (STR, LOF, and HIF) and path Widths (Wide, Narrow) for the data shown in Figs. 3, 5 and 6, including: step errors, means (*μ*) and variability (σ) for Margin of Stability (*MoS*_*L*_), and Probability of Instability (*PoI*_*L*_). ANOVA results (F-statistics and p-values) are provided for main effects of Frequency, Width, and Frequency×Width. In cases where Frequency×Width interactions were significant, we considered main effects results (marked ‘p*’) to be unreliable. We drew conclusions instead from the Tukey’s pairwise comparisons, as these account for the effects of the interactions. Significant differences are indicated in bold.

| Data In | Variable | Frequency | Width | Frequency × Width | Tukey’s (Frequency Effects) | Tukey’s (Width Effects) |
| --- | --- | --- | --- | --- | --- | --- |
| Fig. 3: | $\ln(\text{Step Errors}+0.001)$ | $F_{(2,259)} = 113.10$<br>$p^* < 0.001$ | $F_{(1,259)} = 535.39$<br>$p^* < 0.001$ | $F_{(2,259)} = 11.37$<br>$p < 0.001$ | STRW-LOFW: $p = 0.987$<br>**STRW-HIFW: $p < 0.001$**<br>**LOFW-HIFW: $p < 0.001$**<br>**STRN-LOFN: $p < 0.001$**<br>**STRN-HIFN: $p < 0.001$**<br>**LOFN-HIFN: $p < 0.001$** | **STRW-STRN: $p < 0.001$**<br>**LOFW-LOFN: $p < 0.001$**<br>**HIFW-HIFN: $p < 0.001$** |
| Fig. 5A: | $\mu(MoS_L)$ | $F_{(2,261)} = 2,016.96$<br>$p^* < 0.001$ | $F_{(1,261)} = 403.57$<br>$p^* < 0.001$ | $F_{(2,259)} = 46.07$<br>$p < 0.001$ | STRW-LOFW: $p = 0.232$<br>**STRW-HIFW: $p < 0.001$**<br>**LOFW-HIFW: $p < 0.001$**<br>**STRN-LOFN: $p < 0.001$**<br>**STRN-HIFN: $p < 0.001$**<br>**LOFN-HIFN: $p < 0.001$** | **STRW-STRN: $p < 0.001$**<br>**LOFW-LOFN: $p < 0.001$**<br>**HIFW-HIFN: $p < 0.001$** |
| Fig. 5B: | $\ln(\sigma(MoS_L))$ | $F_{(2,259)} = 6,747.67$<br>$p^* < 0.001$ | $F_{(1,259)} = 6.92$<br>$p^* < 0.01$ | $F_{(2,259)} = 15.24$<br>$p < 0.001$ | **STRW-LOFW: $p < 0.001$**<br>**STRW-HIFW: $p < 0.001$**<br>**LOFW-HIFW: $p < 0.001$**<br>**STRN-LOFN: $p < 0.001$**<br>**STRN-HIFN: $p < 0.001$**<br>**LOFN-HIFN: $p < 0.001$** | STRW-STRN: $p = 0.736$<br>LOFW-LOFN: $p = 1.000$<br>**HIFW-HIFN: $p < 0.001$** |
| Fig. 6: | $\ln(PoI_L+0.001)$ | $F_{(2,259)} = 1093.58$<br>$p^* < 0.001$ | $F_{(1,259)} = 60.09$<br>$p^* < 0.001$ | $F_{(2,259)} = 11.90$<br>$p < 0.001$ | **STRW-LOFW: $p < 0.001$**<br>**STRW-HIFW: $p < 0.001$**<br>**LOFW-HIFW: $p < 0.001$**<br>**STRN-LOFN: $p < 0.001$**<br>**STRN-HIFN: $p < 0.001$**<br>**LOFN-HIFN: $p < 0.001$** | **STRW-STRN: $p < 0.001$**<br>**LOFW-LOFN: $p < 0.001$**<br>**HIFW-HIFN: $p = 0.985$** |

**Figure 3.**
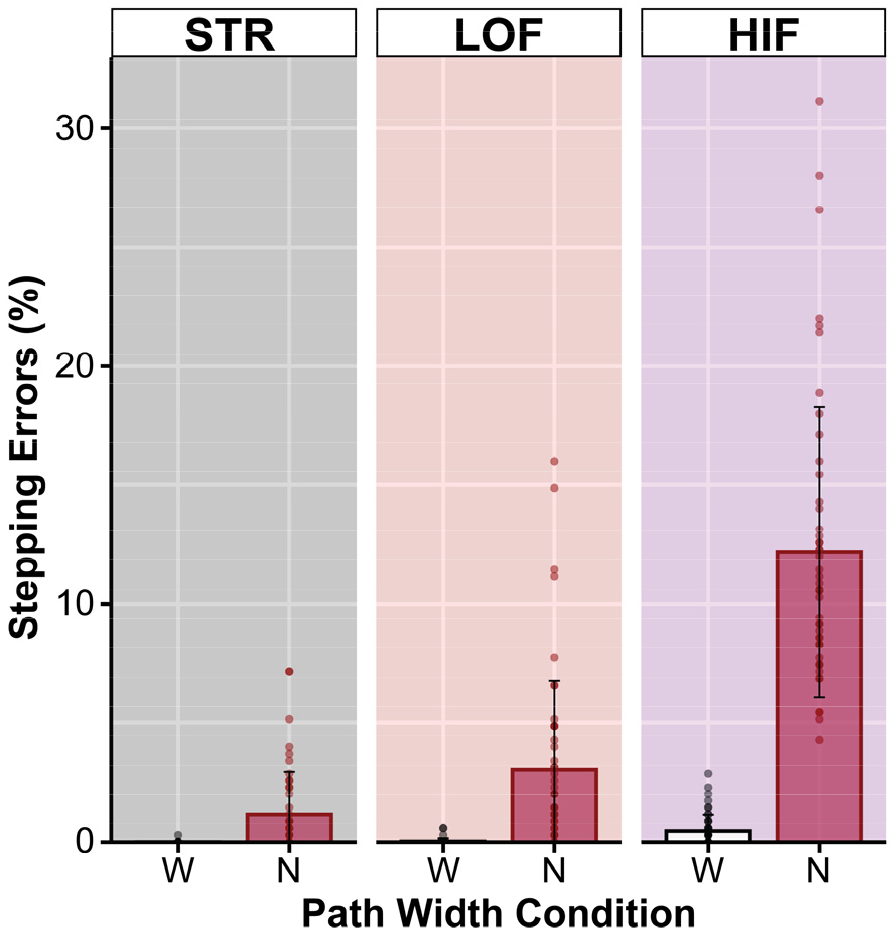
Percent stepping errors on the wide (W) and narrow (N) paths for each path oscillation frequency: STR (grey), LOF (red), and HIF (purple). Bar plots represent the mean values. The overlaid markers are individual data points for each trial for each participant. Results of statistical analyses are detailed in Table 2.

### Lateral Margins of Stability

Qualitatively, step-to-step time series of *MoS*_*Lmin*_ exhibited increasingly larger-amplitude fluctuations from the STR paths to LOF paths, and much larger still from the LOF paths to HIF paths (Fig. 4).

**Figure 4.**
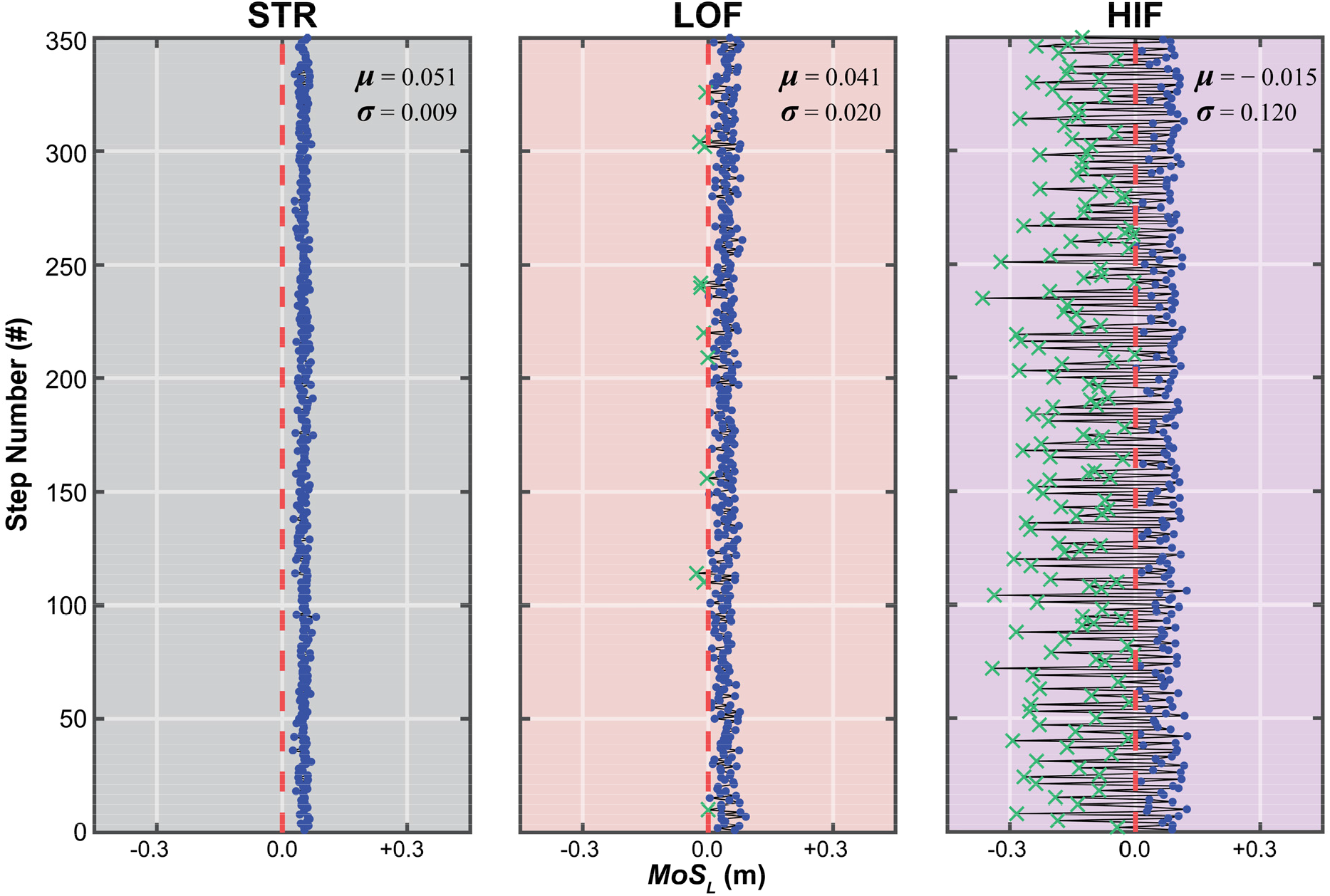
Example minimum lateral Margin of Stability (*MoS*_*Lmin*_) time series (350 consecutive steps each) for three representative trials from a typical participant walking on a Wide path for each path oscillation frequency: STR, LOF, and HIF. The vertical red dashed line indicates *MoS*_*Lmin*_ = 0. Solid blue markers (•) indicate steps where *MoS*_*Lmin*_ > 0, which were considered stable. Green markers (×) indicate steps where *MoS*_*Lmin*_ < 0, which were considered unstable. The within-trial means (*μ*) and standard deviations (*σ*) of each *MoS*_*Lmin*_ time series are shown on each corresponding subplot.

People exhibited smaller means, *μ*(*MoS*_*Lmin*_), on paths of narrower width (p < 0.001), and on narrow paths of increased oscillation frequency (all p < 0.001; Fig. 5A; Table 2). Across wide paths, people exhibited similar *μ*(*MoS*_*Lmin*_) on STR and LOF (p = 0.232), but smaller *μ*(*MoS*_*Lmin*_) on HIF (both p < 0.001). People exhibited greater variance, *σ*(*MoS*_*Lmin*_), on the Narrow vs. the Wide HIF path (p < 0.001; Fig. 5B; Table 2), but similar variance for both path widths for both STR (p = 0.736) and LOF (p = 1.000). People also exhibited increased *σ*(*MoS*_*Lmin*_) with increased path sinuosity (from STR to LOF to HIF), regardless of path width (all p < 0.001; Fig. 5B; Table 2).

**Figure 5.**
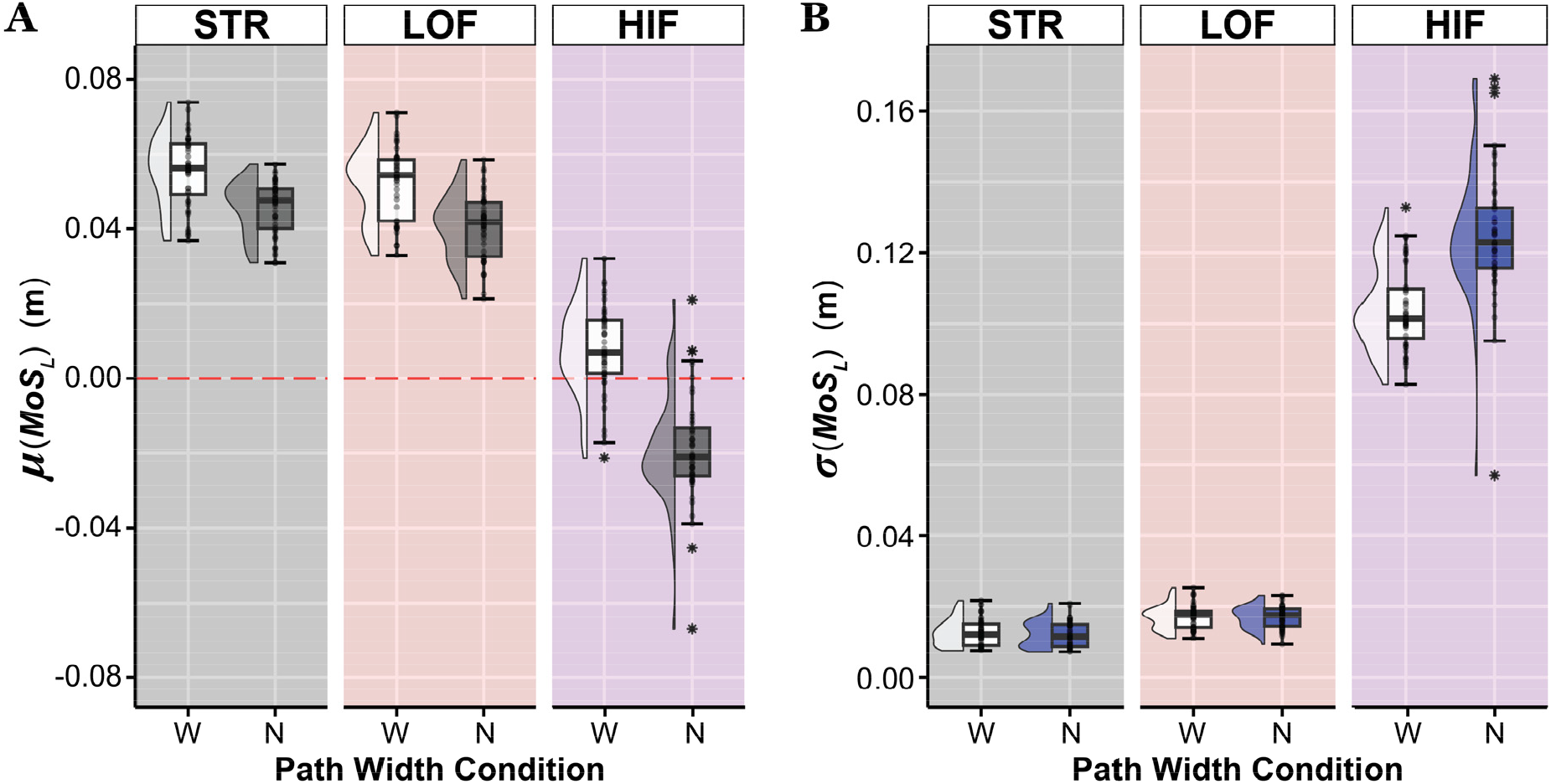
**(A)** Within-trial means (*μ*) of minimum lateral Margin of Stability (*MoS*_*Lmin*_) on the wide (W) and narrow (N) paths for each path oscillation frequency: STR (grey), LOF (red), and HIF (purple). **(B)** Corresponding within-trial step-to-step standard deviations (*σ*) of *MoS*_*Lmin*_ for each path combination. Box plots show the medians, 1^st^ and 3^rd^ quartiles, and whiskers extending to 1.5 × interquartile range. Values beyond this range are shown as individual asterisks. The overlaid markers are individual data points for each participant from two separate trials. Half-violin plots are also shown for visual reference. The red horizontal dashed line in (**A**) indicates *MoS*_*Lmin*_ = 0. Results of statistical analyses are detailed in Table 2.

### Lateral Probability of Instability

For each trial, that trial’s mean, *μ*(*MoS*_*Lmin*_) (Fig. 5A), and standard deviation, *σ*(*MoS*_*Lmin*_) (Fig. 5B), were used to compute that trial’s *PoI*_*L*_ (Eq. 4) (Fig. 6). People exhibited higher *PoI*_*L*_ (p < 0.001) on the narrower compared to wider paths for both STR and LOF (Fig. 6; Table 2), but similar *PoI*_*L*_ at both path widths for HIF (p = 0.985).

**Figure 6.**
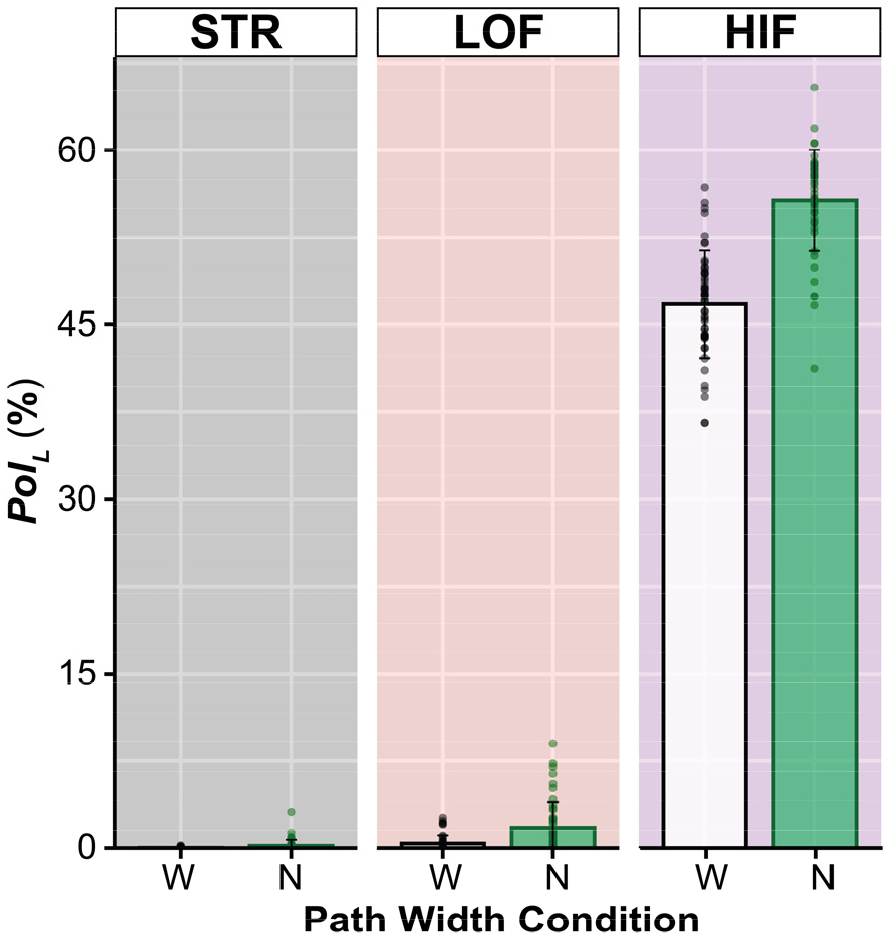
Lateral Probability of Instability (*PoI*_*L*_) on the wide (W) and narrow (N) paths for each path oscillation frequency: STR (grey), LOF (red), and HIF (purple). Bar plots represent the mean values. The overlaid markers are individual data points for each participant from two separate trials. Results of statistical analyses are detailed in Table 2.

For both path widths individually, people exhibited very low *PoI*_*L*_ on the STR paths, slightly but significantly (p < 0.001) higher *PoI*_*L*_ on the LOF paths, and *much* higher *PoI*_*L*_ on the HIF paths (all p < 0.001) (Fig. 6). Pooled across both path widths, participants’ mean probability of taking unstable steps on the STR paths was only *PoI*_*L*_ ≈ 0.11% (range [0%, 3.15%]). This equates to taking 1 unstable step every ∼909 steps (range [∞, ∼32] steps). On the LOF paths, these increased to mean *PoI*_*L*_ ≈ 1.08% (range [0%, 8.99%]), or 1 unstable step every ∼93 (range [∞, ∼11]) steps. On the HIF paths, these increased to mean *PoI*_*L*_ ≈ 51.2% (range [∼36.6%, ∼65.4%]), or 1 unstable step every ∼2.0 (range [∼2.7, ∼1.5]) steps. Thus, the HIF paths in particular *substantially* disrupted people’s ability to maintain stable walking (e.g., as seen in Fig. 4).

## DISCUSSION

Everyday walking often entails navigating crowded and/or non-straight paths (Bergsma et al., 2021; Degond et al., 2013; Glaister et al., 2007). These tasks require people to adapt their stepping to precisely maneuver (Acasio et al., 2017; Wu et al., 2015). However, this maneuverability can increase one’s risk of experiencing instability (Desmet et al., 2024; Tillman et al., 2022). Many older adults fall because they cannot adequately negotiate such non-straight walking tasks (Robinovitch et al., 2013). Despite their prevalence, we know little about how path width and/or shape affect lateral stability during walking. This study therefore examined real-world-like walking on winding paths of varying sinuosity and width. The results reveal how such tasks challenge people’s ability to maintain lateral balance while walking.

Our first hypothesis, that people would take more unstable steps when walking on *narrow* paths was supported. On the Narrow paths, participants made more stepping errors for each path frequency (Fig. 3), walked with smaller mean *MoS*_*Lmin*_ on all three path types (Fig. 5A), more variable *MoS*_*Lmin*_ on HIF paths (Fig. 5B), and exhibited statistically increased instability risk on the Narrow STR and LOF paths (Fig. 6). Our second hypothesis, that people would take more unstable steps on windier paths was very strongly supported.

Participants nearly always made increasingly more stepping errors from STR to LOF to HIF paths (Fig. 3), walked with both smaller mean *MoS*_*Lmin*_ (Fig. 5A) and more variable *MoS*_*Lmin*_ (Fig. 5B) from STR to LOF to HIF paths, and most importantly, exhibited significantly increased instability risk from STR to LOF to HIF paths (Fig. 6). Our third hypothesis that simultaneously narrow and windier paths would further exacerbate participant’s ability to maintain balance was partly supported. People did not make more stepping errors with increased path sinuosity on Wide paths from STR to LOF, but did on HIF and Narrow paths (Fig. 3). For *σ*(*MoS*_*Lmin*_), path width differences not observed for STR and LOF paths, became significant (Table 2) on HIF (Fig. 5B) paths.

On the Narrow paths, across all sinuosity conditions, people walked with significantly smaller mean *MoS*_*Lmin*_ (Fig. 5A; Table 2). This was most likely because they needed to adopt narrower step widths (Arvin et al., 2016; Kazanski et al., 2024; McAndrew Young and Dingwell, 2012; Schrager et al., 2008). However, despite these smaller *μ*(*MoS*_*Lmin*_), which would typically be interpreted as indicating greater instability (Watson et al., 2021), we observed minimal corresponding changes in people’s probability to take unstable steps (Fig. 6). From the Wide to Narrow, participants exhibited significant (p < 0.001), but modest increases in *PoI*_*L*_, from 0.025% to 0.20% (i.e., from taking an unstable step once every ∼4,000 steps to once every ∼500 steps) on STR paths, and from 0.39% to 1.78% (i.e., from taking an unstable step once every ∼256 steps to once every ∼56 steps) on LOF paths. However, the increases observed in *PoI*_*L*_ from the Wide to Narrow paths for HIF (Fig. 6) were not statistically significant (p = 0.985; Table 2). These results further support prior findings (Kazanski et al., 2022, 2024) that measures of *μ*(*MoS*_*Lmin*_), despite their prevalence (Watson et al., 2021), do not quantify risk of taking unstable steps, whereas *PoI*_*L*_ does (Fig. 2C).

On the winding paths (LOF & HIF), participants took more unstable (*MoS*_*Lmin*_ < 0) steps, especially on the HIF paths, where participants took unstable steps as often as once every ∼1-3 steps (Figs. 4 & 6). By comparison, in a prior experiment, we subjected young and older adults to unpredictable lateral physical perturbations (Kazanski et al., 2022). Those perturbations only induced, on average, *PoI*_*L*_ < ∼3% (range [0%, ∼21%]), or 1 unstable step every ∼33 (range [∞, ∼5]) steps ((Kazanski et al., 2022); Fig. 8B). Hence, walking on winding paths appears potentially far more laterally destabilizing than physically perturbing people. This may seem somewhat counterintuitive given that, in the current experiment, participants experienced *no* external perturbations and were given full visual information about their path well in advance to plan their foot placements (Matthis et al., 2017; Matthis et al., 2018). However, unlike in (Kazanski et al., 2022) where participants could step where they chose, participants here were instructed to keep their feet *on their paths*. This constrained where they could take their next step, which most likely accounts for their increased instability (Bruijn and van Dieën, 2018).

For HIF trials, *MoS*_*Lmin*_ time series (Fig. 4) appeared potentially skewed. Since Eq. (4) estimates *PoI*_*L*_ assuming a Gaussian distribution (Fig. 2C), we computed the skewness of each *MoS*_*Lmin*_ timeseries. We then also calculated the actual percentage of unstable *MoS*_*Lmin*_ < 0 steps taken within each trial, and correlated these against our *PoI*_*L*_ estimates. While STR and LOF trials were mostly minimally skewed, most HIF trials were moderately negatively skewed (Fig. 7A). Correspondingly, *PoI*_*L*_ predicted % unstable steps well for both STR and LOF, but somewhat overestimated % unstable steps for HIF (Fig. 7B). Hence, assuming normality somewhat limited the accuracy of our *PoI*_*L*_ predictions, but only for the HIF trials.

**Figure 7.**
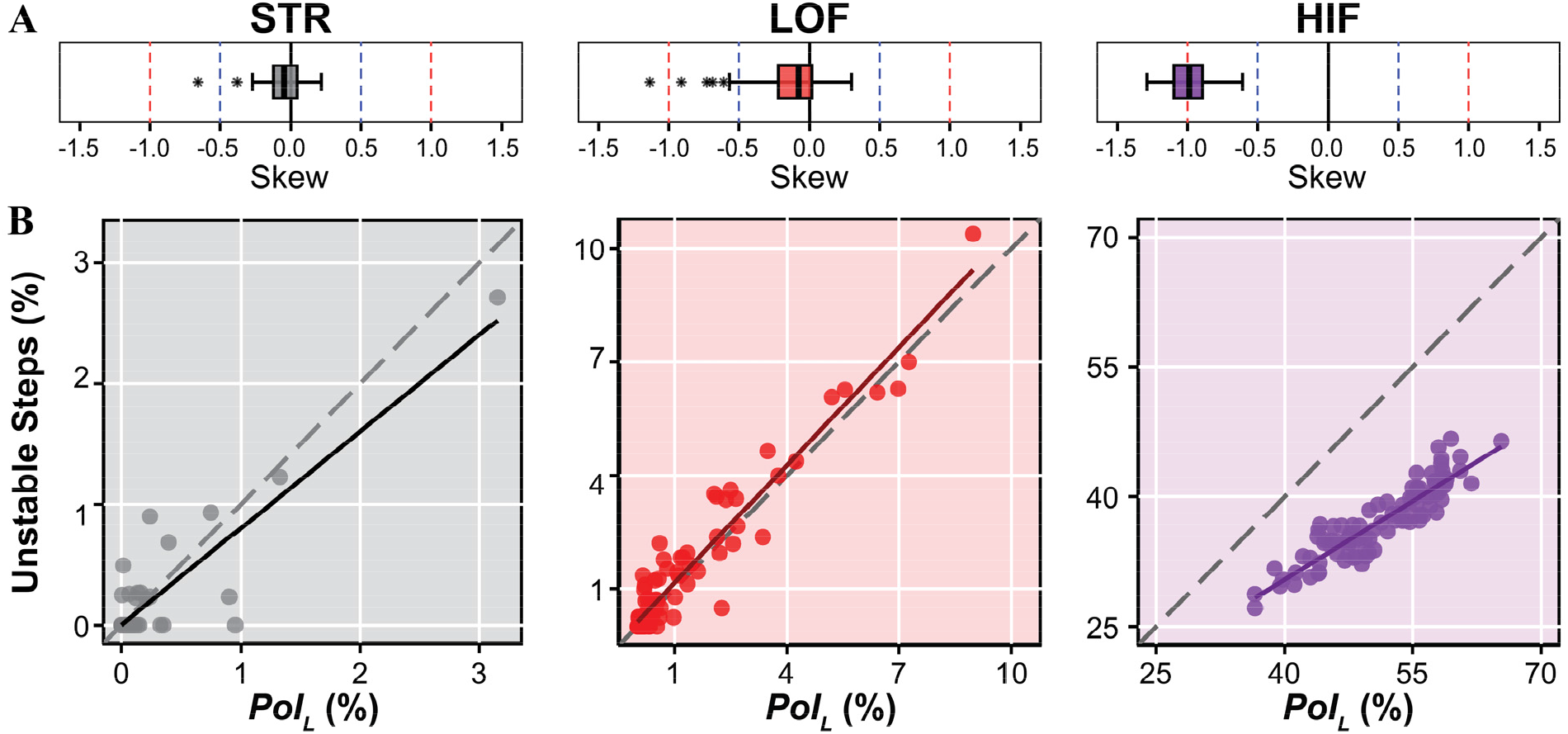
**A)** Boxplots of skewness values computed from *MoS*_*Lmin*_ time series (e.g., Fig. 5) computed from all trials pooled across both path Widths for each of the STR, LOF, and HIF path oscillation frequency conditions. Step-to-step *MoS*_*Lmin*_ distributions were predominantly symmetrical (−0.5 < skewness < +0.5) for STR and LOF trials, but mostly moderately negatively skewed (−1.0 < skewness < −0.5) for HIF. **B)** Correlations of *PoI*_*L*_ to direct measurements of percent unstable steps taken for each path oscillation frequency. The diagonal grey dashed line indicates the identity line. *PoI*_*L*_ predicts % Unstable Steps very well for STR and LOF paths, but over-estimates % Unstable Steps for HIF.

Importantly however, these differences in no way change our conclusions here, given the very large *PoI*_*L*_ for HIF compared to STR and LOF (Fig. 6). Future studies should check for skewness in their *MoS*_*Lmin*_ data before estimating *PoI*_*L*_. In some contexts, it might be more appropriate to directly calculate the percent of unstable (*MoS*_*Lmin*_ < 0) steps taken, such as when sample sizes are large (i.e., many steps per trial) and/or instability risk is high (e.g., as in HIF trials here). However, in contexts where long trials are not feasible and/or instability risk is low/moderate (e.g., as in STR or LOF trials here), one may not observe enough unstable steps to get good direct estimates of the actual percent of *MoS*_*Lmin*_ < 0 steps taken, and Eq. (4) may be preferrable. Alternatively, one could modify Eq. (4) to incorporate skewness in addition to mean and standard deviation.

In their recent review article, Jain et al. pointed out that metrics like *MoS*_*L*_ “…fail to capture the probabilistic characteristics of behavior associated with risk” (Jain et al., 2024). This is particularly true when researchers simply compute average values of *MoS*_*L*_ across multiple steps. Conversely, *PoI*_*L*_ explicitly predicts a person’s *probability* to experience an unstable (i.e., *MoS*_*L*_ < 0) step over some sequence of steps, thus directly resolving the deficiency highlighted by (Jain et al., 2024). Additionally, however, when one does take a step with *MoS*_*L*_ < 0, they also then need to rectify that instability on subsequent step(s). *PoI*_*L*_ does not, by itself, address how people might do this, but other time series analyses can (Desmet et al., 2022; Dingwell and Cusumano, 2019).

Moreover, because *PoI*_*L*_ is merely another statistical metric computed from the same *MoS*_*L*_ distribution used to compute means and standard deviations, *PoI*_*L*_ can be applied to a wide range of laboratory (e.g., (Ochs et al., 2021; Onushko et al., 2019)) and/or real-world (e.g., (Tillman et al., 2022)) tasks, and also patient populations (e.g., (Watson et al., 2021)), just as measures of *μ*(*MoS*_*L*_) have been. In particular, winding (non-straight) walking tasks occur in many real-world contexts (Bergsma et al., 2021; Degond et al., 2013; Glaister et al., 2007) and can be particularly hazardous for those with impaired gait (Robinovitch et al., 2013). Because *PoI*_*L*_ is conceptually consistent with Hof’s definition (Hof, 2008; Hof et al., 2005), whereas *μ*(*MoS*_*L*_) is not (Kazanski et al., 2022), *PoI*_*L*_ is a far more relevant metric to compute for those impaired persons and/or in those varied walking contexts.

In this experiment, despite participants’ clearly taking numerous unstable steps (Fig. 4 & 6) on the winding paths, *no participant fell*. Taking a step with *MoS*_*Ln*_ < 0 does not necessitate failure (Gill et al., 2019) – but instead, only that one must use strategies to regain balance that are *not* available to an inverted pendulum (Hof et al., 2005; Kazanski et al., 2022). The primary strategy is to take a *step* (Redfern and Schumann, 1994; Townsend, 1985).

Even simple bipeds generally can choose among infinite stepping options to minimally remain “viable” (i.e., “not fall”) (Patil et al., 2022; Zaytsev et al., 2018), and switching stepping goals at each step can, in principle, allow them to walk effectively forever (Patil et al., 2022). Humans use such strategies when they make discrete maneuvers (Desmet et al., 2022). Moreover, viability gives rise to “*semi*stability”, which allows expansive versatility of walking that is both viable and purposeful (Patil et al., 2024). That participants here took so many unstable steps (e.g., Fig. 4) without falling strongly suggests people can employ such highly-versatile stepping strategies to continuously trade off stability for maneuverability (Acasio et al., 2017) at each step.

Here, continuously winding paths compromised participant’s stability, demonstrating how people trade off stability for maneuverability. The increased likelihood for people taking an unstable step on these winding paths emphasizes the importance of individuals being able to frequently and successfully enact mechanisms to regain their balance and mitigate falls in destabilizing contexts. To improve everyday mobility in at-risk populations, interventions can target strategies, such as stepping adjustments or weight shifting, that enable people to quickly and effectively respond to balance disturbances.

## ACKNOWLEDGEMENTS

The authors thank Dr. David M. Desmet and Dr. Meghan E. Kazanski for their contributions and technical support throughout data collections. This work was supported by NIH grants 1-R01-AG049735 & 1-R21-AG053470 to JBD & JPC.

## REFERENCES

[Dataset] Render, A.C., Cusumano, J.P., Dingwell, J.B., 2024. Data: Healthy Human Adults Walking on Winding Paths, Dryad Digital Repository (10.5061/dryad.3tx95×6rb), 1 ed.

Acasio, J., Wu, M.M., Fey, N.P., Gordon, K.E., 2017. Stability-maneuverability trade-offs during lateral steps. Gait Posture 52, 171–177.

Arvin, M., Mazaheri, M., Hoozemans, M.J.M., Pijnappels, M., Burger, B.J., Verschueren, S.M.P., van Dieën, J.H., 2016. Effects of narrow base gait on mediolateral balance control in young and older adults. J. Biomech. 49, 1264–1267.

Bergsma, B., Hulleman, D.N., Wiedemeijer, M.M., Otten, E., 2021. Foot placement variables of pedestrians in community setting during curve walking. Gait Posture 86, 120–124.

Bohnsack-McLagan, N.K., Cusumano, J.P., Dingwell, J.B., 2016. Adaptability of Stride-To-Stride Control of Stepping Movements in Human Walking. J. Biomech. 49, 229–237.

Bruijn, S.M., van Dieën, J.H., 2018. Control of human gait stability through foot placement. J. R. Soc. Interface 15, 1–11.

Curtze, C., Buurke, T.J.W., McCrum, C., 2024. Notes on the Margin of Stability. J. Biomech. 166, 112045.

da Silva Costa, A.A., Moraes, R., Hortobágyi, T., Sawers, A., 2020. Older adults reduce the complexity and efficiency of neuromuscular control to preserve walking balance. Experimental Gerontology 140, 111050.

Dean, J.C., Alexander, N.B., Kuo, A.D., 2007. The effect of lateral stabilization on walking in young and old adults. IEEE Trans. Biomed. Eng. 54, 1919–1926.

Degond, P., Appert-Rolland, C., Moussaïd, M., Pettré, J., Theraulaz, G., 2013. A Hierarchy of Heuristic-Based Models of Crowd Dynamics. J Stat Phys 152, 1033–1068.

Desmet, D.M., Cusumano, J.P., Dingwell, J.B., 2022. Adaptive Multi-Objective Control Explains How Humans Make Lateral Maneuvers While Walking. PLoS Comput. Biol. 18, e1010035.

Desmet, D.M., Kazanski, M.E., Cusumano, J.P., Dingwell, J.B., 2024. How Healthy Older Adults Enact Lateral Maneuvers While Walking. Gait Posture 108, 117–123.

Dingwell, J.B., Cusumano, J.P., 2019. Humans Use Multi-Objective Control to Regulate Lateral Foot Placement When Walking. PLoS Comput. Biol. 15, e1006850.

Dingwell, J.B., Render, A.C., Desmet, D.M., Cusumano, J.P., 2023. Generalizing Stepping Concepts to Non-Straight Walking. J. Biomech. 161, 111840.

Gill, L., Huntley, A.H., Mansfield, A., 2019. Does the margin of stability measure predict medio-lateral stability of gait with a constrained-width base of support? J. Biomech. 95, 109317.

Glaister, B.C., Bernatz, G.C., Klute, G.K., Orendurff, M.S., 2007. Video task analysis of turning during activities of daily living. Gait Posture 25, 289–294.

Hak, L., Houdijk, H., Steenbrink, F., Mert, A., van der Wurff, P., Beek, P.J., van Dieën, J.H., 2012. Speeding up or slowing down?: Gait adaptations to preserve gait stability in response to balance perturbations. Gait Posture 36, 260–264.

Hak, L., Houdijk, H., Steenbrink, F., Mert, A., van der Wurff, P., Beek, P.J., van Dieën, J.H., 2013. Stepping strategies for regulating gait adaptability and stability. J. Biomech. 46, 905–911.

Havens, K.L., Mukherjee, T., Finley, J.M., 2018. Analysis of biases in dynamic margins of stability introduced by the use of simplified center of mass estimates during walking and turning. Gait Posture 59, 162–167.

He, C., Xu, R., Zhao, M., Guo, Y., Jiang, S., He, F., Ming, D., 2018. Dynamic stability and spatiotemporal parameters during turning in healthy young adults. BioMedical Engineering OnLine 17, 127.

Ho, T.K., Kreter, N., Jensen, C.B., Fino, P.C., 2023. The choice of reference frame alters interpretations of turning gait and stability. J. Biomech. 151, 111544.

Hof, A.L., 2008. The ‘extrapolated center of mass’ concept suggests a simple control of balance in walking. Hum. Mov. Sci. 27, 112–125.

Hof, A.L., Gazendam, M.G.J., Sinke, W.E., 2005. The condition for dynamic stability. J. Biomech. 38, 1–8.

Hurt, C.P., Grabiner, M.D., 2015. Age-related differences in the maintenance of frontal plane dynamic stability while stepping to targets. J. Biomech. 48, 592–597.

Jain, S., Schweighofer, N., Finley, J.M., 2024. Aberrant decision-making as a risk factor for falls in aging. Frontiers in Aging Neuroscience 16, 1384242.

Kazanski, M.E., Cusumano, J.P., Dingwell, J.B., 2022. Rethinking Margin of Stability: Incorporating Step-to-Step Regulation to Resolve the Paradox. J. Biomech. 144, 111334.

Kazanski, M.E., Cusumano, J.P., Dingwell, J.B., 2023. How Older Adults Regulate Lateral Stepping on Narrowing Walking Paths. J. Biomech. 160, 111836.

Kazanski, M.E., Cusumano, J.P., Dingwell, J.B., 2024. How Older Adults Maintain Lateral Balance While Walking on Narrowing Paths. Gait Posture 113, 32–39.

Matthis, J.S., Barton, S.L., Fajen, B.R., 2017. The critical phase for visual control of human walking over complex terrain. Proc. Natl. Acad. Sci. USA 114, E6720–E6729.

Matthis, J.S., Yates, J.L., Hayhoe, M.M., 2018. Gaze and the Control of Foot Placement When Walking in Natural Terrain. Curr. Biol. 28, 1224-1233.e1225.

McAndrew Young, P.M., Dingwell, J.B., 2012. Voluntary changes in step width and step length during human walking affect dynamic margins of stability. Gait Posture 36, 219–224.

McAndrew Young, P.M., Wilken, J.M., Dingwell, J.B., 2012. Dynamic Margins of Stability During Human Walking in Destabilizing Environments. J. Biomech. 45, 1053–1059.

Moussaïd, M., Helbing, D., Theraulaz, G., 2011. How simple rules determine pedestrian behavior and crowd disasters. Proc. Natl. Acad. Sci. USA 108, 6884–6888.

Musselman, K.E., Yang, J.F., 2007. Walking tasks encountered by urban-dwelling adults and persons with incomplete spinal cord injuries. J. Rehabil. Med. 39, 567–574.

Ochs, W.L., Woodward, J., Cornwell, T., Gordon, K.E., 2021. Meaningful measurements of maneuvers: People with incomplete spinal cord injury ‘step up’ to the challenges of altered stability requirements. J. Neuroeng. Rehabil. 18, 46.

Onushko, T., Boerger, T., Van Dehy, J., Schmit, B.D., 2019. Dynamic stability and stepping strategies of young healthy adults walking on an oscillating treadmill. PLoS ONE 14, e0212207.

Parkkari, J., Kannus, P., Palvanen, M., Natri, A., Vainio, J., Aho, H., Vuori, I., Järvinen, M., 1999. Majority of Hip Fractures Occur as a Result of a Fall and Impact on the Greater Trochanter of the Femur: A Prospective Controlled Hip Fracture Study with 206 Consecutive Patients. Calcif Tissue Int 65, 183–187.

Patil, N.S., Dingwell, J.B., Cusumano, J.P., 2022. Viability, Task Switching, and Fall Avoidance of the Simplest Dynamic Walker. Scientific Reports 12, 8993.

Patil, N.S., Dingwell, J.B., Cusumano, J.P., 2024. A model of task-level human stepping regulation yields semistable walking. J. R. Soc. Interface In Press, (Pre-Print: 10.1101/2024.1103.1105.583616).

Patla, A.E., Frank, J.S., Winter, D.A., Rietdyk, S., Prentice, S., Prasad, S., 1993. Age-related changes in balance control system: initiation of stepping. Clin. Biomech. 8, 179–184.

Redfern, M.S., Schumann, T., 1994. A model of foot placement during gait. J. Biomech. 27, 1339–1346.

Robinovitch, S.N., Feldman, F., Yang, Y., Schonnop, R., Leung, P.M., Sarraf, T., Sims-Gould, J., Loughin, M., 2013. Video capture of the circumstances of falls in elderly people residing in long-term care: an observational study. Lancet 381, 47–54.

Schrager, M.A., Kelly, V.E., Price, R., Ferrucci, L., Shumway-Cook, A., 2008. The effects of age on medio-lateral stability during normal and narrow base walking. Gait Posture 28, 466–471.

Tillman, M., Molino, J., Zaferiou, A.M., 2022. Frontal plane balance during pre-planned and late-cued 90 degree turns while walking. J. Biomech. 141, 111206.

Townsend, M.A., 1985. Biped Gait Stabilization Via Foot Placement. J. Biomech. 18, 21–38.

Twardzik, E., Duchowny, K., Gallagher, A., Alexander, N., Strasburg, D., Colabianchi, N., Clarke, P., 2019. What features of the built environment matter most for mobility? Using wearable sensors to capture real-time outdoor environment demand on gait performance. Gait Posture 68, 437–442.

van Dieën, J.H., Pijnappels, M., Bobbert, M.F., 2005. Age-related intrinsic limitations in preventing a trip and regaining balance after a trip. Safety Science 43, 437–453.

Watson, F., Fino, P.C., Thornton, M., Heracleous, C., Loureiro, R., Leong, J.J.H., 2021. Use of the margin of stability to quantify stability in pathologic gait – a qualitative systematic review. BMC Musculoskeletal Disorders 22, 597.

Wu, M., Matsubara, J.H., Gordon, K.E., 2015. General and Specific Strategies Used to Facilitate Locomotor Maneuvers. PLoS ONE 10, e0132707.

Wu, M.M., Brown, G., Gordon, K.E., 2017. Control of Locomotor Stability in Stabilizing and Destabilizing Environments. Gait Posture 55, 191–198.

Yang, Y., Komisar, V., Shishov, N., Lo, B., Korall, A.M.B., Feldman, F., Robinovitch, S.N., 2020. The Effect of Fall Biomechanics on Risk for Hip Fracture in Older Adults: A Cohort Study of Video-Captured Falls in Long-Term Care. Journal of Bone and Mineral Research 35, 1914–1922.

Zaytsev, P., Wolfslag, W., Ruina, A., 2018. The Boundaries of Walking Stability: Viability and Controllability of Simple Models. IEEE Trans Robot. 34, 336–352.

Zeni, J.A., Richards, J.G., Higginson, J.S., 2008. Two simple methods for determining gait events during treadmill and overground walking using kinematic data. Gait Posture 27, 710–714.

